# Efficient inhibition of fusion inhibitor HY3000 peptide to SARS-CoV-2 emerging EG.5, EG.5.1 and BA.2.86 variants

**DOI:** 10.1101/2023.09.28.559747

**Authors:** Lili Wu, Anqi Zheng, Yangming Tang, Xiaoyun Wang, Yue Gao, Wenwen Lei, Guizhen Wu, Qihui Wang, George Fu Gao

## Abstract

SARS-CoV-2 continues to evolve and spread. Recently, the Omicron EG.5 lineage, bearing an additional F456L mutation in spike (S) protein compared to its ancestor XBB.1.9.2, and its sub-variant EG.5.1, which carries a further Q52H mutation, have raised concerns due to their increased prevalence and extended immune escape properties. Additionally, an alarming variant, BA.2.86, has also garnered global concern because it contains over 30 amino acid mutations in its S protein compared to BA.2, including more than 10 changes in receptor-binding domain (RBD), reminiscent of the appearance of the Omicron variant in late 2021. Therefore, there is an urgent need to assess the effectiveness of current vaccines and therapeutics against EG.5, EG.5.1 and BA.2.86. In our previous work, we reported the design and broad-spectrum antiviral activity of a peptide fusion inhibitor HY3000 against SARS-CoV-2 and its variants including XBB.1.5. Here, we continued to evaluate the inhibitory potency of the HY3000 peptide against the prevailing EG.5 and EG.5.1, as well as XBB.1.16, FL.1.5.1, FY.3 and BA.2.86. Our data indicated that the peptide retained its potent inhibitory activities against these variants, indicating its potential as a good virus fusion inhibitor with broad-spectrum therapeutic effect against current and future SARS-CoV-2 variants. Currently, the HY3000 has been finished in Phase II clinical trial in China and has also been approved to conduct clinical investigation by U.S. Food and Drug Administration (FDA), suggesting a good application prospect against the ongoing COVID-19.

## Main text

Severe acute respiratory syndrome coronavirus 2 (SARS-CoV-2) continues to pose a significant threat to the world, as it continually evolves and gives rise to multiple variants and sub-variants. Recently, the Omicron EG.5 linage, which was first detected in Indonesia on 17 February 2023, has raised concerns due to its increased prevalence and extended immune escape properties, according to the risk analysis by the World Health Organization (WHO)^1^. As of 26 September 2023, EG.5 and its sub-linages have been reported in 77 countries with shared 40,099 genome sequences in GISAID database (https://gisaid.org/hcov19-variants/). EG.5 has become the dominant strain in the United States according to Centers for Disease Control and Prevention (CDC), accounting for 24.5% of SARS-CoV-2 infections (https://covid.cdc.gov/covid-data-tracker/#variant-proportions), while EG.5.1 is the dominant strain in China on the past three months (https://nmdc.cn/ncovn/china/statistics). Additionally, an alarming linage BA.2.86, with numerous mutations in its spike (S) protein related to BA.2 or XBB.1.5, has been detected in 22 countries (https://gisaid.org/hcov19-variants/), which is reminiscent of the appearance of the Omicron variant in late 2021 and has raised significant concerns^2^. WHO has designated it as a variant under monitoring (VUM) on 17 August 2023 (https://www.who.int/activities/tracking-SARS-CoV-2-variants). Given the increased prevalence, there is an urgent need to assess the effectiveness of current vaccines and therapeutics against EG.5, EG.5.1 and BA.2.86, as well as to develop broad-spectrum countermeasures.

The EG.5 linage is a descendant of XBB.1.9.2, which shares the same S amino acid profile as XBB.1.5 and XBB.1.9.1. EG.5 carries an additional F456L mutation in the receptor-binding domain (RBD) of the S protein compared to XBB.1.9.2/XBB.1.9.1/XBB.1.5 (**Fig. 1a**). EG.5.1 has an additional Q52H mutation in the N-terminal domain (NTD) of the S protein compared to EG.5. Additionally, FY.3, an another prevailing strain in China, has an additional Y200C mutation in the NTD but lacks F456L compared to EG.5. XBB.1.16 and FL.1.5.1, especially FL.1.5.1, a descendant of XBB.1.9.1, is currently the second dominant strain in the United States, which possesses two additional mutations (F456L and A701V) and a distinct mutation (T478R) in the S protein compared to XBB.1.9.1. However, in contrast to the abovementioned sub-variants with limited mutations, BA.2.86 bears 34 mutations in the S protein with 14 mutations in the RBD compared to its putative ancestor BA.2 and 35 mutations in the S protein with 11 mutations in the RBD compared to XBB.1.5, which imply enhanced immune evasion. Notably, all these variants share the same S2 region including heptad repeat 1 (HR1) and HR2, except FL.1.5.1 containing an A701V mutation and BA.2.86 carrying a S939F mutation in HR1 and a P1143L mutation in stem-helix. Thus, we speculate that therapeutics targeting the S2 region, like peptide fusion inhibitor HY3000, which derives from HR2 and targets HR1 of SARS-CoV-2 S (Fig. 1d)^3^, should remain effective against these variants, but need to be experimentally verified.

**Figure 1.**
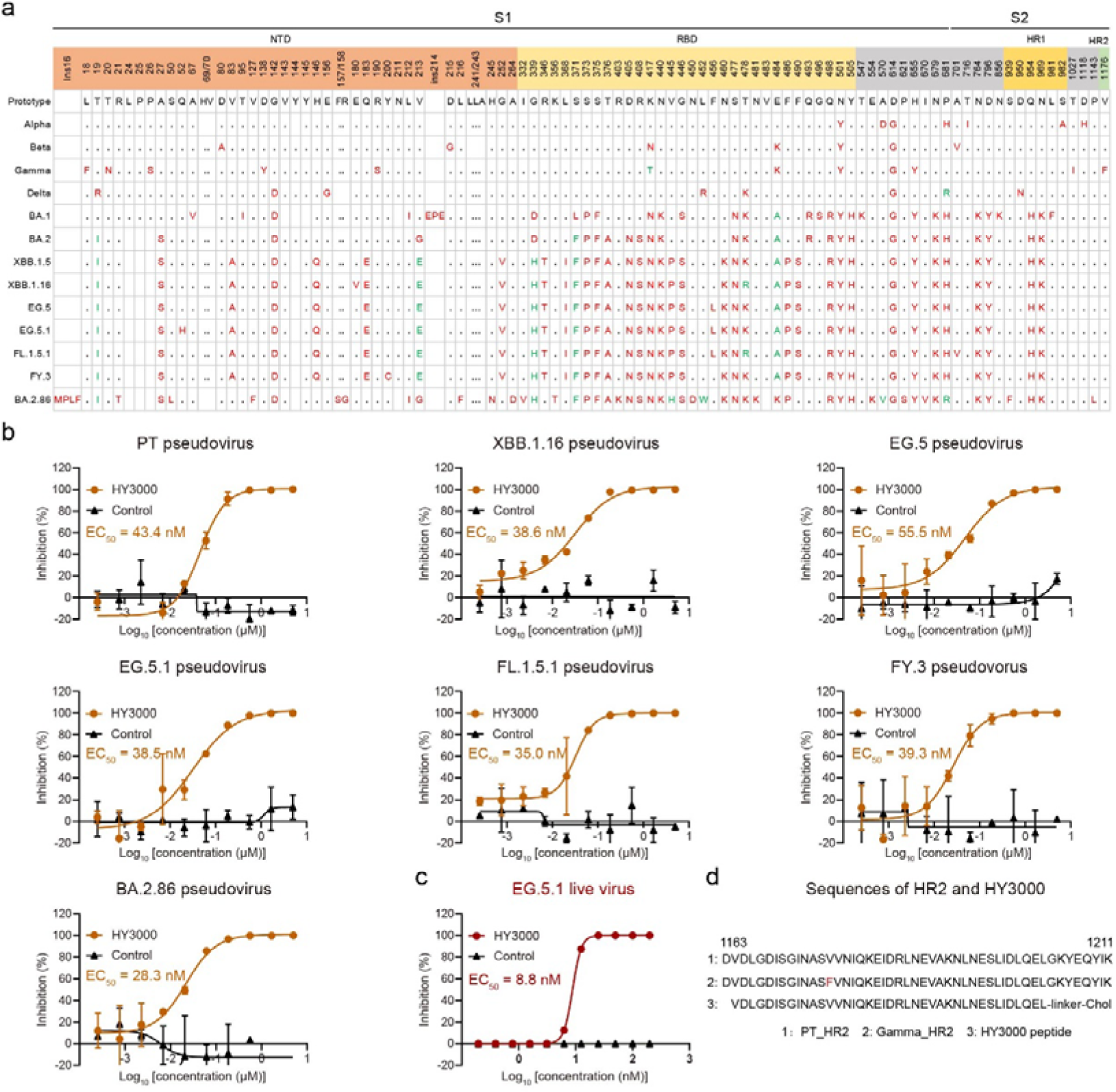
Amino acid mutation mapping of spike (S) proteins from SARS-CoV-2 EG.5, EG.5.1 and BA.2.86, in comparison with major existed variants, and inhibitory potencies of HY3000 peptide against these pseudoviruses and EG.5.1 live virus. **a** Amino acid mutation mapping of S proteins from SARS-CoV-2 EG.5, EG.5.1 and BA.2.86. Compered to SARS-CoV-2 prototype S, substituted amino acids were colored in red, while the presence of a second mutation at the same site was colored in green. Amino acid deletions were marked with blank square. **b** The inhibitory activities of the HY3000 peptide against the SARS-CoV-2 PT, XBB.1.16, EG.5, EG.5.1, FL.1.5.1, FY.3 and BA.2.86 pseudoviruses were evaluated in 293T cells with high expression of the human ACE2 receptor (293T-hACE2). The assays were performed independently twice with two replicates (*n*=2) in each experiment. The EC50 values were the means of two independent experiments, and shown curves were one representative results of two independent experiments. Control, peptide derived from influenza virus. **c** The inhibitory potency of the HY3000 peptide against live EG.5.1 virus was assessed in Vero cells based on the cytopathic effect (CPE) with eight replicates (*n*=8) in the experiment. **d** Sequences of HR2 from SARS-CoV-2 PT and Gamma containing a mutation, as well as HY3000 peptide, were shown.

Previous studies have demonstrated that peptide fusion inhibitors derived from HR2 of the SARS-CoV-2 S protein, as well as glycoproteins of other class I enveloped viruses, exhibited efficient antiviral activities against SARS-CoV-2 and related viruses^3-8^. At the beginning of the COVID-19 pandemic, we designed a peptide fusion inhibitor P3 against SARS-CoV-2, based on our experience in developing peptide inhibitors for SARS-CoV and MERS-CoV^4,9,10^. To improve the inhibitory activity of P3, we further designed a lipopeptide HY3000, which exhibited significantly increased inhibitory activities against SARS-CoV-2 and all tested variants including XBB.1.5/XBB.1.9.1/XBB.1.9.2, and has currently been finished in Phase II clinical trial in China^3^. Because of the excellent antiviral potency, we here further evaluated the inhibitory potency of the HY3000 peptide against the prevailing EG.5 and EG.5.1, as well as XBB.1.16, FL.1.5.1, FY.3 and the alarming BA.2.86.

Using our previous assessment methods, HY3000 peptide showed expectedly comparable inhibitory potencies against EG.5 and EG.5.1 pseudoviruses compared to its activity against the SARS-CoV-2 prototype (PT) strain, with half-maximal effective concentration (EC_50_) values of ∼50 nM (**Fig. 1b**), consistent with our previous results^3^. Moreover, HY3000 peptide also displayed similar inhibitory activities against the pseudotyped XBB.1.16, FL.1.5.1 and FY.3 strains as it against PT, EG.5 and EG.5.1. Notably, HY3000 exhibited slightly higher inhibition against BA.2.86 than it against the above sub-variants, with an EC50 value of 28.3 nM. This result possibly attributed to the S393F mutation in the HR1 of the BA.2.86 strain. When modeling S939 to F939 in the HR1 and HR2 complex, we observed that although the aromatic side chain of F939 was oriented away from HR2, it increased hydrophobic interaction with A1190, V1189 and L1186 in HR2 (**Fig. S1**), thus resulting in the slight improvement. Moreover, HY3000 exhibited potent inhibition against live EG.5.1 strain, with an EC50 of 8.8 nM (**Fig. 1c**), which was similar to its activities against SARS-CoV-2 PT and previous Omicron sub-variants including BA.1, BA.2 and BA.4^3^. These results suggested that the HY3000 peptide fusion inhibitor is a good virus fusion inhibitor with broad-spectrum therapeutic effect against current and future potential SARS-CoV-2 variants and sub-variants. However, due to the limited resources of the Animal Biosafety Level 3 facility, we have not yet evaluated the inhibitory potency of the HY3000 peptide against live EG.5 *in vivo*. Nevertheless, based on our previous studies and the efficacy of the peptide *in vitro*, we speculate that the HY3000 can effectively inhibit live virus infections *in vivo*. Additionally, given the ongoing evolution of SARS-CoV-2, continuous surveillance of variants and continuous evaluation of antiviral effects of the HY3000 peptide inhibitor are necessary in the future.

## Supporting information

This supplemental file contains materials and methods and Supplemental Figure S1.

## Acknowledgments

This work was supported by the National Key R&D Program of China (2023YFC0871300 and 2022YFC2303403) and the National Natural Science Foundation of China (82225021). Q.W. is supported by the Chinese Academy of Sciences (YSBR-010 and Y2022037).

## Author contributions

G.F.G. and Q.W. initiated and coordinated the project. With the help of X.W. and Y.G., A.Z. performed the pseudovirus-based inhibitory assays. W.L. and G.W. evaluated the potency of HY3000 against live virus. Y.T. synthesized the peptide. L.W., Q.W. and G.F.G. wrote the manuscript.

