## Supplementary material for "Efficient inhibition of fusion inhibitor HY3000 peptide to SARS-CoV-2 emerging EG.5, EG.5.1 and BA.2.86 variants": This supplemental file contains materials and methods and Supplemental Figure S1.

**Supplementary information**

**Figure S1**

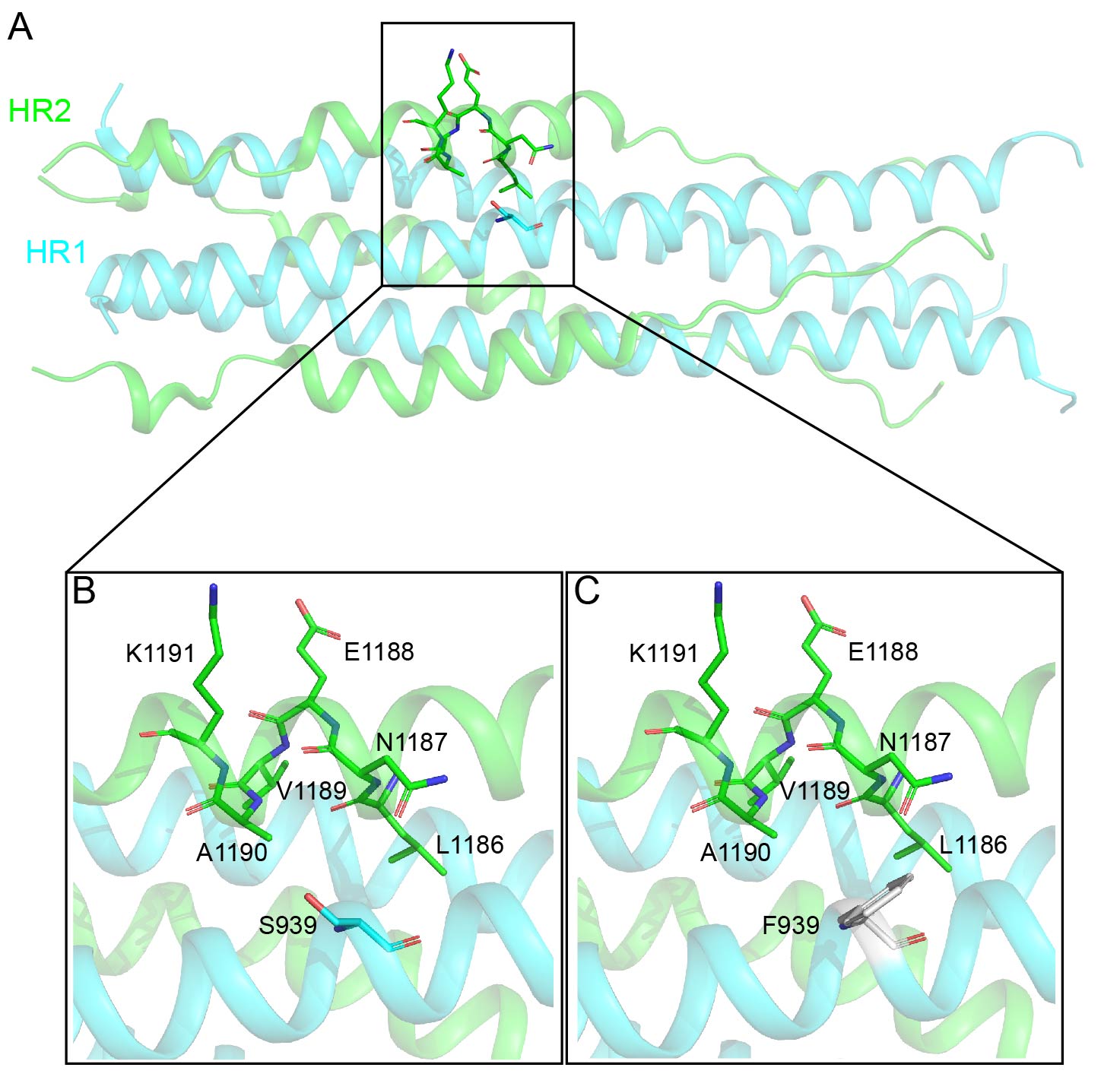

**Figure S1 Structural comparison of the S939F mutation in HR1 and potential residues involved interaction in HR2.** **A** The six-helix bundle structure generated by symmetry operation of the HR1 and HR2 complex (PDB: 6M1V). HR1 and HR2 were colored cyan and green, respectively. S939 in HR1 and involved residues in HR2 were shown as sticks and was further amplified in **B** and **C**. **B** S939 in HR1 and potential residues involved in HR2. **C** Modeled F939 (gray) in HR1 and potential residues involved in HR2.

**Materials and methods**

**Cells**

HEK293T (ATCC, CRL-3216), BHK-21 (ATCC, CCL-10), Vero (ATCC, CCL-81) and HEK293T-hACE2 (Genewiz®) were cultured at 37 ℃ in Dulbecco’s modified Eagle’s medium (DMEM) supplemented with 10% fetal bovine serum (FBS), 1‰ penicillin and 1‰ streptomycin.

**Virus**

The SARS-CoV-2 EG.5.1 strain was isolated and propagated in the National Institute for Viral Disease Control and Prevention, Chinese Center for Disease Control and Prevention.

**Pseudovirus preparation**

The VSV-ΔG-GFP-based SARS-CoV-2 pseudoviruses were prepared as previously described^1^. Briefly, sequences encoding the spike proteins of tested viruses, with 18 amino acid truncations at the C-terminus, were synthesized and cloned into the pCAGGS vector, respectively. The plasmids were transfected into HEK293T cells. After 24 h of transfection, the VSV-ΔG-G-GFP pseudovirus was added to the cells. Following a 2 h of incubation, the medium was removed and replaced with fresh DMEM containing 10 μg/mL of anti-VSV-G antibody (I1‐Hybridoma ATCC® CRL-2700™). After 30 h, the supernatant was collected, centrifuged at 3,000 rpm for 10 min, and filtered through a 0.45 μm filter to remove cell debris. The pseudoviruses were then frozen at -80 ℃ until use. HEK293T-hACE2 cells were used to determine pseudovirus titers, and BHK-21 cells were used to detect residual VSV-ΔG-G-GFP pseudovirus.

**Pseudovirus inhibition assay**

HKE293T-hACE2 cells were seeded in 96-well plates 24 h before pseudovirus infection. Peptides were serially diluted 2- or 3-fold with DMEM containing 2% FBS, and 50 μL of the diluted peptides were mixed with each pseudovirus in equal volumes. The mixture was then incubated at 37 ℃ for 1 h and subsequently added to the cells. After 15 h, the number of cells expressing GFP was quantified using a CQ1 confocal image cytometer (Yokogawa). The inhibitory activities of the peptide were analyzed using GraphPad Prism 8.

**Live SARS-CoV-2 virus inhibition assay**

The inhibitory activity of peptide against live SARS-CoV-2 EG.5.1 virus was determined based on the cytopathic effect (CPE). Briefly, 50 μL of 2-fold serial dilutions (starting from 800 nM) of peptide were incubated with an equal volume of 100 TCID_50_ of virus at 37°C for 1 h. Then, 100 μL of suspended Vero cells (2×10^4^ cells/well) were added to the mixtures and incubated for 4 days at 37°C. CPE was observed and recorded. All experiments were performed in the Biosafety Level 3 (BSL-3) facility of National Institute for Viral Disease Control and Prevention, Chinese Center for Disease Control and Prevention. The data were analyzed using GraphPad Prism 8.

**Reference**

1	Wu, L. *et al.* A pan-coronavirus peptide inhibitor prevents SARS-CoV-2 infection in mice by intranasal delivery. *Sci China Life Sci (in press)*.
